# No solid evidence of soil carbon loss under warming in tropical forests along a 3000 m elevation gradient

**DOI:** 10.1101/2020.02.10.941310

**Authors:** Zhongkui Luo, Xiaowei Guo, Osbert Jianxin Sun

## Abstract

Soil organic carbon (SOC) decomposition is inherently sensitive to temperature. As such, a big concerning is the potential SOC loss under climatic warming, but field empirical evidences are lacking, particularly in tropical forest soils in which ∼10% of global SOC is stored. Recently Nottingham et al. (2019) assessed the data collected from a novel experiment translocating soils across a 3000 m tropical forest elevation gradient to mimic temperature changes *in situ*, and concluded that warming caused considerable SOC loss. However, this conclusion was based on a metric with a strong assumption that soil cores translocated to other elevations on average had the same initial SOC content to control soil cores reinstalled at their original elevation. Because of limited replicates (*n* =3) in the data, an approach ignoring spatial heterogeneity of SOC content may undermine the credibility of the results. Here, we used a nonparametric bootstrap approach to re-analyze the data, explicitly taking data variability into account. Contrary to Nottingham et al. (2019), we found that SOC content did not show significant differences among translocated soils from the same elevation origin. Further looking into six chemical fractions determined by ^13^C NMR spectroscopy shown that they had similar, insignificant response to translocation-induced temperature changes, which also does not support the conclusion of Nottingham et al (2019) that labile SOC is more sensitive to warming. We concluded that temperature changes did not significantly alter either total SOC content or its six chemical fractions after five years of shift of temperature regimes in tropical forests. This may largely due to thermal adaptation of microbial decomposition and environmental constrains (e.g., low pH) which suppress the effect of temperature changes. Longer term experiment with more sampling replicates are required to maximize the value of soil translocation experiments to address the effect of warming on SOC dynamics.

## Introduction

Soil organic carbon (SOC) pool in tropical forest soils accounts for ∼40% of SOC stock in global forests (Pan et al., 2011) and ∼10% in global soils (Batjes, 2016; Le Quere et al., 2016). As the inherent temperature sensitivity of SOC decomposition (Davidson & Janssens, 2006), it is vital to understand how SOC in tropical forests responds to climatic warming. Earth system models usually predict SOC loss under warming including in tropical forest soils, rising a big concern of positive SOC loss - climatic warming feedbacks (Allison, Wallenstein, & Bradford, 2010). Nevertheless, there are no solid, consistent empirical evidences to prove those model projections. Results from field warming experiments are inconclusive; the effects of warming on SOC balance are observed to be positive, neutral or negative, depending on study-specific ecosystems, experimental manipulation (such as duration), local soil and climatic conditions, and other confounding factors (e.g., Sistla et al., 2013; Pries, Castanha, Porras, & Torn, 2017; and two data syntheses by Crowther et al., 2016 and van Gestel et al., 2018). A 26-year soil warming experiment at Harvard forest (a temperate forest) indeed observed three multiyear phases of soil microbial respiration: from the first phase of decreasing respiration, to the second phase of stable respiration, to the third phase of increasing respiration, due to changes in substrate availability and microbial community functioning (Melillo et al., 2017). In tropics, the data is particularly lacking (Crowther et al., 2016; van Gestel et al., 2018), inhibiting our understanding and quantification of the fate of SOC under climatic warming in tropical areas.

By conducting a field experiment in tropical forests, recently Nottingham et al. (2019) addressed the effects of long-term (5 years) soil warming on SOC content and a series of microbial properties. Intriguingly, to do so, they translocated soils among four tropical forest sites along a 3000 m elevation gradient in Peru to generate an average temperature change of ± 15 °C. After five years of the translocation, they measured SOC content and a suite of other soil chemical and biological properties. By assessing the data (hereafter we call it “Nottingham dataset”), they concluded that “warming caused a considerable loss of soil carbon” (Nottingham et al. 2019). They also measured six chemical fractions including carbonyl (165-190 ppm), O-aryl (140-165 ppm), aryl (110-140 ppm), di-O-alkyl (92-110 ppm), O-alkyl (46-92 ppm), and alkyl (0-46 ppm), using ^13^C NMR spectroscopy. By assessing the fraction data, they concluded that SOC loss was related to the lability of SOC component fractions. Particularly, labile SOC fractions were more sensitive to temperature changes and their loss was the major contributor to total SOC loss.

In Nottingham et al. (2019), however, there was a strong assumption underlying the estimation of the effects of temperature change. That is, soil cores translocated to other elevations on average have the same initial SOC content to control soil cores reinstalled at their original elevation. As there are only three replicates in the data, an approach ignoring spatial heterogeneity of SOC content may over- or under-estimate the effect of warming. Focusing on total SOC content and its chemical component fractions, in this study we reassessed the Nottingham dataset by explicitly taking into account data variability, and found that neither total SOC content nor its six chemical fractions was significantly affected by elevation shift-induced temperature changes.

### Reassessment of Nottingham dataset

In Nottingham et al. (2019), relative response ratios (RR) were calculated to estimate the effect of temperature change (which was represented by elevation shift in Nottingham et al. 2019 due to its close correlation with temperature change, R^2^ = 0.99):

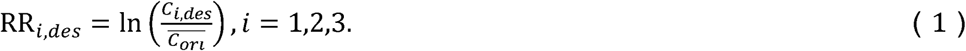

where *c*_*i,des*_ is the variable of interest (i.e., total SOC content and its six chemical component fractions in this study) of the *i*^th^ replicate of soils translocated to other elevation destination, and 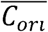 is the average of three replicates of the interested variable in the control soil cores reinstalled at original elevation. As such, three RRs were obtained for each destination of each soil origin (a total of four soil origins from four elevations). Based on these RRs, Nottingham et al (2019) conducted a series of statistical analyses such as regression and ANOVA. However, their analyses ignored the potential effect of data variability on the results.

In this re-analysis, we first conducted pair-wise comparisons of total SOC and its chemical fractions (the data were natural log-transformed before conducting the comparison) among translocated soils of the same origin using ANOVA (pair-wise Tukey post-hoc test, which controls for Type I error). This analysis enables us to directly assess whether or not soil translation has induced significant changes in SOC content and in its chemical fractions. Embracing the benefit of RR for quantifying “the relative effect of translocation (warming or cooling) on each property independently to soil type” (Nottingham et al. 2019), and meanwhile taking into account the potential effect of data variability on results, here we used a non-parametric bootstrap approach to quantify the relationship between RR and temperature changes due to soil translocation along the 3000 m elevation gradient. The non-parametric approach is not only robust to departures from the normal distribution but also explicitly take data variability into account (Fox & Weisberg, 2011). As there are three replicates for both control soils at the origin location and soils translocated to other destinations, we can calculate nine estimations of RR for each destination. Based on this, we conducted 200 bootstrapping simulations. For each simulation, RR was randomly selected from the nine RR for each translocated soil. A linear regression model was fitted treating RR and temperature change (due to soil translocation) as response and independent variables, respectively. Then, we calculated the average of RR for each translocated soil based on the 200 random draws, and a linear regression model was fitted using the average RRs. The significance of all regressions (i.e., a total of 201 regressions including 200 bootstrapping regressions plus one average regression) was tested at *P* < 0.05.

It should be noted that Nottingham et al (2019) focused on the relationship of RR of soil C with elevation shift rather than directly with temperature change, albeit the close correlation between temperature change and elevation shift (R^2^=0.99). As elevation shift may include changes of a series of other ecosystem properties (e.g., radiation, humidity, rainfall regimes although total amount of rainfall entering to the experimental soils was controlled in their study, see discussions below) other than temperature, our reassessment directly used the elevation shift-induced average temperature changes as a predictor variable of RR. The data is available from Nottingham et al. (2019).

### Total SOC changes

The ANOVA results indicated that total SOC content did not show significant difference among translocated soils of the same origin for all four soil origins (Fig. 1). It is apparent that great variability existed for SOC content in some translocated soils of the same origin (Fig. 1). The bootstrapping regression simulations indicated that only 16 of the 200 simulations were significant (*P* < 0.05) for RR of soil C content (Fig. 2). When specifically assessing the relationship of average RR of SOC content with temperature changes, the relationship was also insignificant (*P* = 0.14) with a R^2^ of 0.21 (Fig. 2). In Nottingham et al (2019), a 3.86% significant (*P* < 0.05) decline of SOC content per 1 °C temperature increase was estimated. Based on the result in this reassessment, however, this decline was insignificant and only 1.4% per 1 °C temperature increase (i.e., the regression coefficient for the temperature change). These results demonstrate that five years of translocation did not significantly changed SOC content in soils from the same origin, and is consistent with a global data synthesizing of forest soil respiration in mineral soils (Giardina & Ryan, 2000). Expanding the dataset used in (Crowther et al., 2016), a recent global data synthesis of field warming experiments also indicated that SOC stock does not significantly respond to warming (van Gestel et al., 2018), although no tropical data was included.

**Fig. 1.**
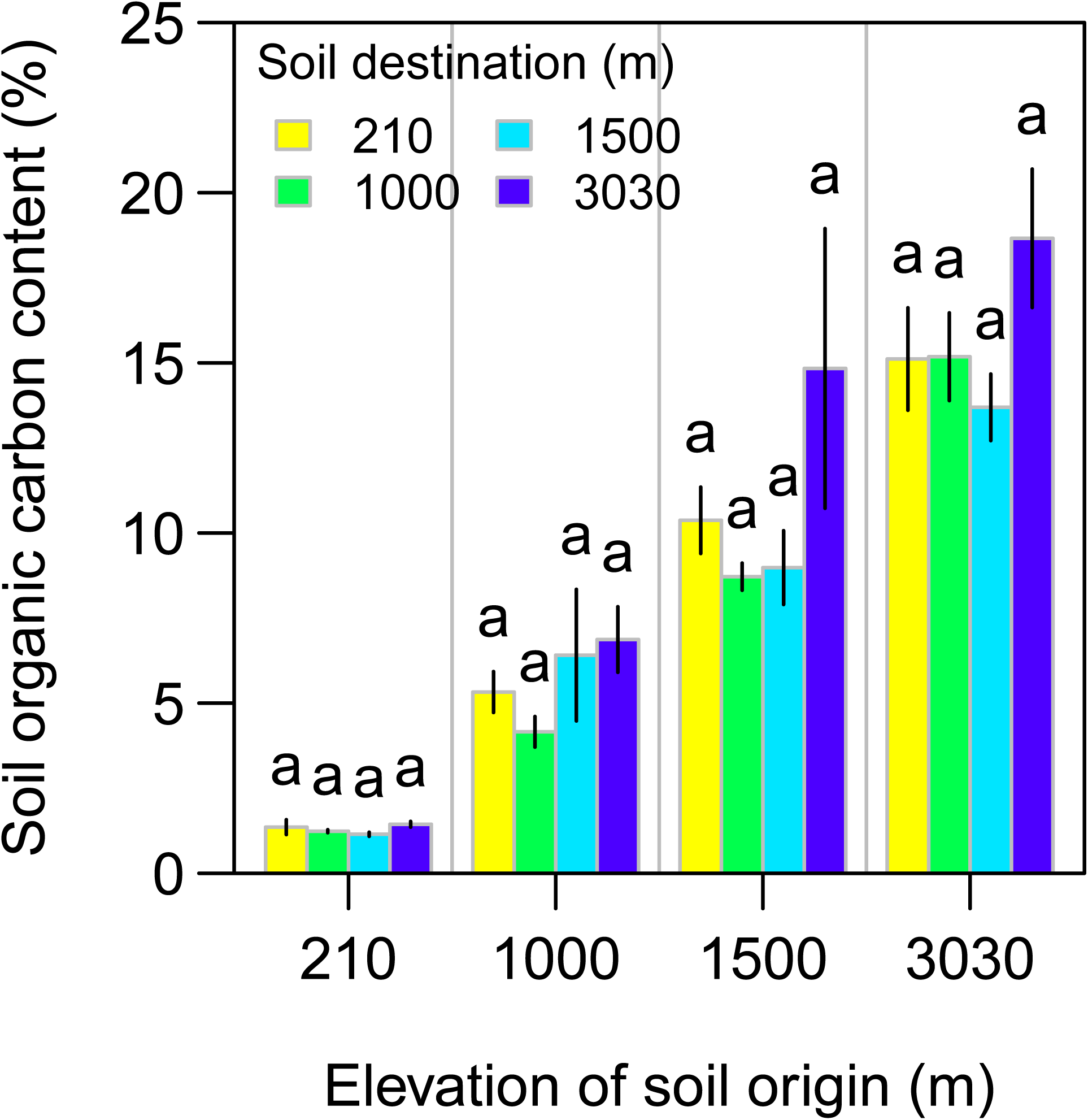
Effects of soil translocation on soil carbon content. Different letters above the bars for the same soil origin group indicate significant difference at *P* < 0.05. Error bars show one standard error. Please note that the increasing pattern just shows that soil carbon increases with elevation and does not relate to the effect of temperature changes.

**Fig. 2.**
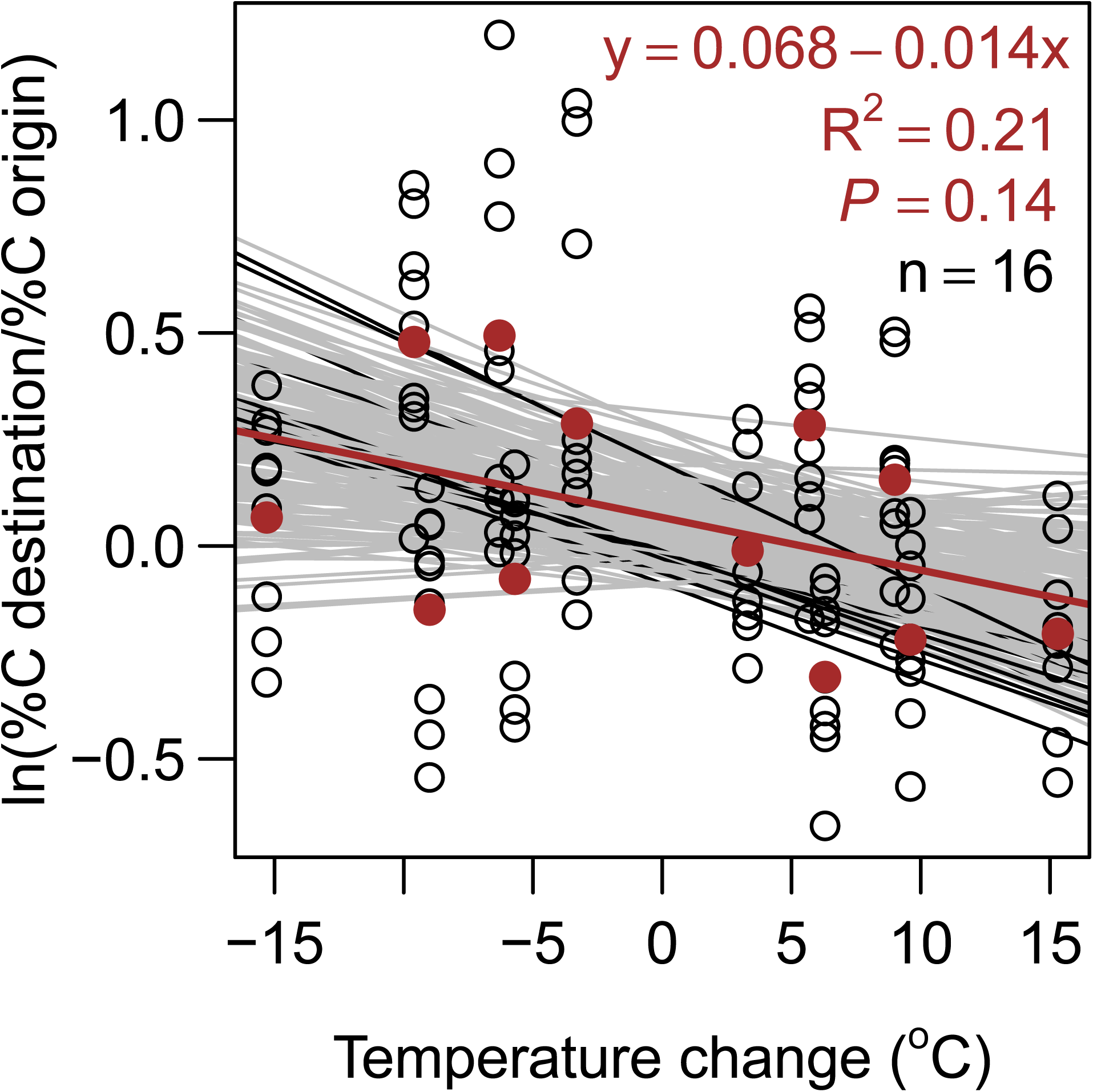
Relationship of the log relative response ratio (RR) of soil carbon content with temperature changes. Circles show nine possible RR taking into account combinations of three replicates at soil destination and origin. Thin lines show 200 bootstrapping regression lines with RR randomly selected from the nine RR values, while black and grey ones indicate that the regression is significant (*P* < 0.05) and insignificant (*P* > 0.05), respectively. *n* shows the number of significant regression lines. Brown solid points are the average of the nine RR values, while brown lines are the corresponding regression lines and brown texts in the plots show the statistics of the regression lines based on average values.

Why is total SOC tolerant towards temperature changes (both warming and cooling)? Although limitations of the data itself (see the discussion below) may result in that the real temperature response of SOC cannot be detected (i.e., Type II error from the perspective of statistics), we discuss the possibility of four mechanisms to explain such persistent SOC content under different temperature regimes: 1) microbial thermal adaptation, 2) substrate depletion, 3) nutrient limitation for microbial carbon acquisition, and 4) environmental constraints such as low pH.

The thermal adaptation hypothesis suggests that microbial community may gradually adapt to warming via adjusting microbial physiology and/or shifting community composition (Bradford, 2013; Luo, Wan, Hui, & Wallace, 2001). Although microbial respiration is sensitive to temperature shift at onset, it will decline towards the pre-warmed/cooled rates over time due to microbial adaptation to temperature shift. A laboratory incubation of three contrasting soils under temperatures ranging from 5 to 25 °C indicated that soil respiration was only significantly different at the start several days of the incubation and gradually reached to a similar rate, regardless of substrate availability and initial microbial community composition (Tang, Sun, Luo, He, & Sun, 2018). For this reason, the long-term effect of warming on total SOC content could be negligible, although significant short-term changes in soil respiration at the start of an experiment. Indeed, microbial measurements after five years of the translocation in Nottingham et al. (2019) indicated that microbial community composition was not significantly different evidenced by the observation that the majority of microbial taxa was unaffected by temperature changes (please see Fig. 2 in Nottingham et al. 2019). If microbes do not adapt to temperature changes, it is reasonable to infer that microbial community composition may be significant different among the translocated soils. Other studies at the same sites also suggested that microbial respiration was adapted to warming (Nottingham, Bååth, Reischke, Salinas, & Meir, 2019).

Besides potential microbial adaptation, microbial activity and functioning could be constrained by other environmental factors (Manzoni, Taylor, Richter, Porporato, & Agren, 2012). The low pH in the studied soils (pH < 4 for all soils, Nottingham et al. 2019) could play a critical role in regulating microbial growth and activity in the studied soils. A number of studies have demonstrated that microbial activity and growth are inhibited in acidic soils (Jones et al., 2009; Rousk, Brookes, & Bååth, 2010). A field study in a silty loam soil at Rothamsted research demonstrated that all microbial variables including fungal and bacterial growth were universally inhibited below pH 4.5 (Rousk, Brookes, & Baath, 2009). It is highly probably that microbial processes are less sensitivity to warming at low pH soils, as pH rather than temperature is the limiting factor of microbial carbon decomposition.

It is less possible for substrate depletion and nutrient limitation for microbial carbon acquisition to take effective in the studied soils. Substrate availability and quality (e.g., the carbon: nitrogen ratio of soil organic matter) are substantially different among soils from different elevations (see Table S1 in Nottingham et al. 2019). If substrate depletion does occur, total SOC as well as its six chemical fractions should be to some extent different in terms of their temperature response in soils from different elevations (i.e., different origin). However, in all soils, none of the six chemical fractions shown significant difference (see results below), although substrate availability is substantially different among the soils. For the same reason, nutrient limitation would be not the reason, as the soils studied from different elevations have distinct nutrient reserve. For example, soil carbon: nitrogen ratios range from 4.07 to 19.17, carbon: phosphorus ratios from 41.52 to 253.71, and resin-extractable phosphorus ranges from less than 0.8 mg kg^−1^ soil to more than 79.99 mg kg^−1^ soil. If nutrient is a limiting factor, temperature response of SOC should present some significant difference among the soils due to the distinct nutrient availability.

### Changes in SOC chemical composition

Fig. 3 shows the ANOVA results of the effect of elevation shift on six chemical component fractions in soils from four elevation origins. Except Carbonyl, O-Aryl and Aryl contents in the soil from the 3300 m elevation at its original elevation were significantly different from that in corresponding soils translocated to other elevations, translocation did not significantly influence the content of all six chemical fractions in all four soils. Large data variability also existed for the chemical fractions, particularly in soils with relatively high SOC content. It is interesting to note that both total SOC content and its six chemical fractions shown greater variability in cooler climate at the 3300 m elevation (Figs. 1 and 3). Bootstrapping regressions on the relationship between temperature changes and the RR of the contents of six chemical fractions indicated that only were 2 of 200 simulations significant for Carbonyl (Fig. 4a) and Aryl (Fig. 4c), 11 for Alkyl (Fig. 4f), 24 for O-Alkyl and Di-Alkyl (Fig. 4d and e), and 26 for O-Aryl (Fig. 4b), demonstrating the importance of quantifying data variability. When assessing the relationship of average RR of SOC chemical fractions with temperature changes, the relationship was only marginal (*P* = 0.086) for O-Aryl with a R^2^ of 0.27 (Fig. 4).

**Fig. 3.**
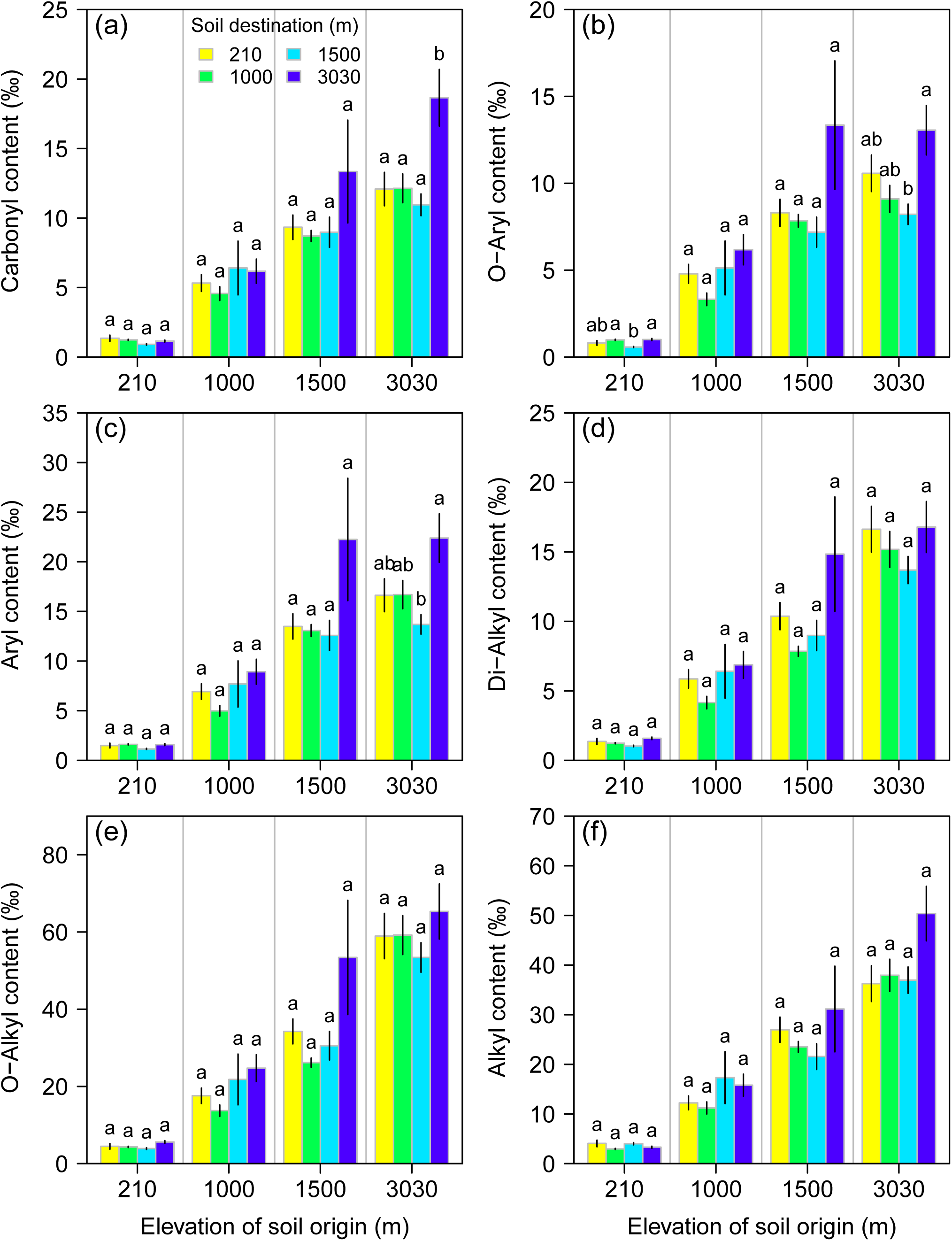
Effects of soil translocation on six chemical fractions of soil carbon. Six chemical fractions are carbonyl (165-190 ppm, a), O-aryl (140-165 ppm, b), aryl (110-140 ppm, c), di-O-alkyl (92-110 ppm, d), O-alkyl (46-92 ppm, e), and alkyl (0-46 ppm, f), determined by ^13^C NMR spectroscopy. See Nottingham et al (2019) for details of the six chemical fractions. Different letters above the bars for the same soil origin group indicate significant difference at P < 0.05. Error bars show one standard error. Please note that the increasing pattern just shows that soil carbon from different elevations are different.

**Fig. 4.**
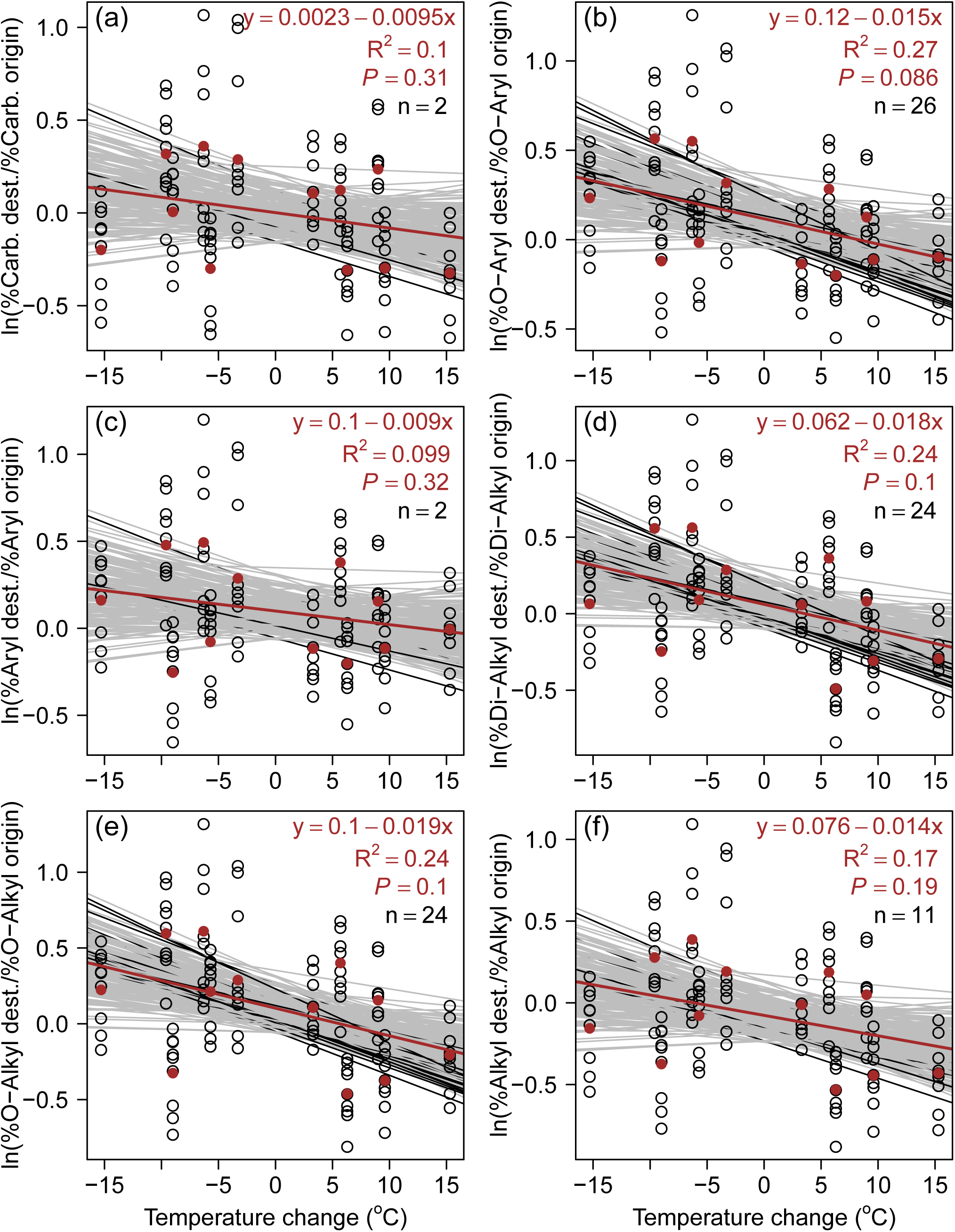
Relationship of log relative response ratio (RR) of six chemical soil carbon fractions with temperature changes. Six chemical fractions are carbonyl (165-190 ppm, a), O-aryl (140-165 ppm, b), aryl (110-140 ppm, c), di-O-alkyl (92-110 ppm, d), O-alkyl (46-92 ppm, e), and alkyl (0-46 ppm, f), determined by ^13^C NMR spectroscopy. See Nottingham et al (2019) for details of the six chemical fractions. Lines, points and legends share the same explanation to that in Fig. 2.

Nottingham et al (2019) interpreted that the detected soil C loss using their approach “primarily originated from labile C pools”. Our re-analysis demonstrated that neither total SOC content nor its chemical fractions was significantly affected by translocation, while their correlations with temperature changes were highly variable and also insignificant on average. Nottingham et al (2019) found that carbonyl content was significantly and positively correlated to temperature changes (Fig. 2 therein), which is difficult to explain (i.e., why carbonyl increases under warming without any external inputs) and opposite to our re-assessment as shown in Fig. 4a. It is clear that all the six chemical fractions were negatively correlated to temperature changes, although none of the correlations was significant at *P* < 0.05 (Fig. 4). The result of the similar temperature response of six chemical fractions is in line with prevailing recognitions.

In a review paper, Dungait, Hopkins, Gregory, & Whitmore (2012) has concluded that SOC turnover is governed by accessibility rather than chemical recalcitrance. A data-model integration study synthesizing global incubation dataset also found that > 90% of SOC is physically protected against microbial decomposition, and the dynamic physical protection process is the limiting step of overall SOC decomposition (Luo et al., 2017). Empirical evidence also shown that the response of SOC decomposition to temperature is constrained by substrate availability to microbial decomposers rather than SOC chemical recalcitrance (Zimmermann, Leifeld, Conen, Bird, & Meir, 2012, Moinet et al. 2018). Using ^14^C techniques, Vaughn & Torn (2019) separated SOC in an Alaska soil into new and old carbon with distinct chemical structures, and found that the two pools shown similar temperature sensitivities. Above all, growing studies have converged on that chemical SOC fractions exert similar temperature sensitivity to decomposition, and physical accessibility rather than chemical structure of SOC is the limiting step. There may be no exception for SOC in tropical forest soils. It will be interesting to identify that whether different chemical fractions are involved in different physical protection processes (i.e., adsorption to and desorption from minerals, occlusion within soil aggregates, and organo-mineral interactions).

### Limitations of Nottingham dataset and future research

Soil translocation experiment is invaluable for mimicking climate change *in situ*, but there are several limitations in the Nottingham dataset. First, as abovementioned, there are only three replicates in the data. Considering the spatial heterogeneity of soil properties including SOC (Garten, Kang, Brice, Schadt, & Zhou, 2007; Stursova, Barta, Santruckova, & Baldrian, 2016; the data in the Appendix of Nottingham *et al.* 2019 can also demonstrate this), more replicates would enable us to provide more accurate estimations of the temperature response of soil C and its chemical components. Second, the experimental design cannot eliminate the effect of soil moisture discrepancies among the translocated soils from the same origin, albeit translocated tubes were capped with reduction collars or expansion funnels to maintain the same rainfall per square meter (Zimmermann, Meir, Bird, Malhi, & Ccahuana, 2010). Nevertheless, absolute rainfall amount is not the only factor influencing soil moisture. Both rainfall regimes (e.g., the time, frequency and intensity of rainfall events) and upward soil water movement may have marked effects on soil moisture including its temporal dynamics. Different rainfall regimes among the destinations may result in complex soil moisture-temperature interactions and relevant consequences on soil carbon decomposition (Rodrigo, Recous, Neel, & Mary, 1997; Zhou, Hui, & Shen, 2014). Upward movement of water through the open bottom of soil tubes (although a 63 µm nylon mesh at the base of the tubes was installed) may be substantial due to capillary action. In order to avoid potential confounding effects of soil moisture, we suggest that both soil moisture and temperature should be monitored over time, making it possible to explicitly separate the effects of soil moisture and temperature changes as well as quantifying their interactions. Third, the duration of the experiment is five years. Considering normal funding cycles and the difficulty to reach the remote areas of the tropical forests, the experiment should be appreciated. However, five years are too short to observe statistically significant trends of soil carbon changes, particularly for recalcitrant pools (if these pools really exist) which usually have residence times of decades or centuries (Luo, Wang, & Wang, 2019; Schmidt et al., 2011). We would suggest to last the experiment as long as possible to detect clear response of SOC to temperature changes.

Another significant confounding factor influencing the results is that plant carbon inputs to the translocated soils were excluded. As pointed out by Nottingham et al (2019), plant carbon inputs to the soil may offset the changes in SOC (although this change is insignificant according to our reassessment) as plant biomass and thus carbon input is generally higher under lower elevation sites where have higher temperature. Except this direct offsetting effect, the absence of carbon inputs has other two kinds of potential consequences on SOC dynamics under temperature changes. First, the absence of carbon input weakens the priming effect. The priming effect is a key process regulating the interaction between new and old SOC (Kuzyakov, 2010; Luo, Wang, & Smith, 2015). Weakening priming effect may have significant consequences on net SOC balance (Rousk, Hill, & Jones, 2015). More importantly, there was no living roots in the soil cores, resulting in the absence of rhizosphere priming effect, which is much more important than the priming effect in the bulk soil as root rhizosphere is a hotspot of microbial growth and activity (Cheng et al., 2014; Zhu et al., 2014). Second, it is unclear whether and how new carbon inputs interact with temperature to affect SOC dynamics. Some evidences from laboratory incubations implied that microbial community structure and functioning respond distinctly to temperature changes under treatments with and without new carbon inputs (e.g., Abro et al., 2011). For these reasons, it should take care when extrapolating results obtained from experiments excluding plant-soil interactions.

## Conclusions

Based on the data collected from translocated soils across a 3000 m tropical forest elevation gradient with ± 15 °C temperature changes, Nottingham et al. (2019) concluded that soil carbon declines 4% per 1 °C warming. Reassessing their data, particularly focusing on measurements of total SOC and its chemical fractions, we did not find significant changes in SOC content, and only found an insignificant decline of ∼1.4% per 1 °C warming. This insignificant response of SOC dynamics to warming may be likely explained by microbial adaptation to temperature changes and/or environmental constrains (such as low pH of the studied soils) which inhibit microbial growth and activity. We also did not find significant changes in six chemical fractions although they potentially reflect distinct chemical recalcitrance, supporting the proposition that chemical SOC fractions may have similar temperature sensitivity, and SOC dynamics are governed by accessibility of substrates to microbial attack rather than the recalcitrance of SOC chemical compounds. Overall, the unique Nottingham dataset fills a gap of data availability in tropical areas, and provides evidence that SOC in tropical soils may be neutral in terms of its response to warming, which is in line with the results in other ecosystems (van Gestel et al., 2018).

## Acknowledgements

We thank the funding support from the National Natural Science Foundation of China (Grant Nos. 31870426, 31470623).

## Authorship

ZL assessed the data and wrote the manuscript. XG and OJS contributed to interpretation and writing.

